# Recording neural reward signals in the real-world using mobile-EEG and augmented reality

**DOI:** 10.1101/2023.08.31.555757

**Authors:** Jaleesa Stringfellow, Omer Liran, Mei-Heng Lin, Travis E. Baker

## Abstract

The electrophysiological response to rewards recorded during laboratory-based tasks has been well documented over the past two decades, yet little is known about the neural response patterns in ‘real-world’ settings. To address this issue, we combined a mobile-EEG system with an augmented reality headset (which blends high definition “holograms” within the real-world) to record event-related brain potentials (ERP) while participants navigated an operant chamber to find rewards. 25 participants (age = 18-43, Male=6, Female=19) were asked to choose between two floating holograms marking a west or east goal-location in a large room, and once participants reached the goal location, the hologram would turn into a reward (5 cents) or no-reward (0 cents) cue. Following the feedback cue, participants were required to return to a hologram marking the start location, and once standing in it, a 3 second counter hologram would initiate the next trial. This sequence was repeated until participants completed 200 trials. Consistent with previous research, reward feedback evoked the reward positivity, an ERP component believed to index the sensitivity of the anterior cingulate cortex to reward prediction error signals. The reward positivity peaked around 235ms post-feedback with a maximal at channel FCz (M=-2.60μV, SD=1.73μV) and was significantly different than zero (p < 0.01). At a behavioral level, participants took approximately 3.38 seconds to reach the goal-location and exhibited a general lose-shift (68.3% ± 3.5) response strategy and were slightly slower to return to the start location following negative feedback (2.43 sec) compared to positive feedback (2.38 sec), evidence of post-error slowing. Overall, these findings provide the first evidence that combining mobile-EEG with augmented reality technology is a feasible solution to enhance the ecological validity of human electrophysiological studies of goal-directed behavior and a step towards a new era of human cognitive neuroscience research that blurs the line between laboratory and reality.

## Introduction

The ability to utilize reward information to adaptively guide behavior to meet current and future goals is essential to successfully navigate through a busy day. Extensive theoretical and empirical work based on simplistic lab-based experiments indicate that goal-directed behavior is largely mediated by key neural targets of the mesocorticolimbic reward system (e.g., orbitofrontal cortex, ventral striatum, prefrontal cortex and anterior midcingulate cortex (ACC), and a dopaminergic teaching signal tethered to prediction of reward outcomes during trial-and-error learning (i.e., reward predication error signals, RPEs) (Schultz, 1998a; Schultz, 1998b, 2002, 2011b; Sutton and Barto, 1998). Current thinking holds that midbrain dopamine neurons distribute information about rewarding events such that phasic bursts and dips in dopamine activity are elicited when events are, respectively, ‘better than expected’ (positive RPE) and ‘worse than expected’ (negative RPE) (Schultz, 1998a, c, 2011a). RPEs are particularly evoked in situations that require continual coordination of perceptions, internal states, and planning of behaviors needed to perform a given task, and allow the mesocorticolimbic reward system to detect rewards, predict future rewards, and use reward information to select and motivate behavior towards a goal (Garrison, Erdeniz and Done, 2013; Niv, Duff and Dayan, 2005; Penner and Mizumori, 2012). Although the neural circuit involved in RPE-related processes has been well defined in simplified and controlled conditions of laboratory settings, it’s unclear how accurately these processes translate to more complex, real-world situations.

To address this issue, the goal of this study was to test a novel mobile-EEG and augmented reality (AR) paradigm aimed to evoke and record RPE-related neural activity during real-world goal-directed behavior. To accomplish this goal, we focused on the role of the ACC in goal-directed behavior and the application of AR to achieve ecological validity in an experimental task setting. Foremost, a prevailing hypothesis holds that the ACC utilizes RPEs to learn the value of rewards for the purpose of selecting and motivating the execution of goaldirected behavior(Holroyd and Coles, 2002; Holroyd and McClure, 2015; Holroyd and Umemoto, 2016; Holroyd and Yeung, 2012). In humans, the reward processing function of ACC can be investigated using a component of the event-related brain potential (ERP) called the reward positivity (also called the feedback-related negativity; for reviews see (Sambrook and Goslin, 2015; Walsh and Anderson, 2012). The reward positivity is observed as a differential response in the ERP to positive and negative feedback received during goal-orientated decision-making tasks (Baker and Holroyd, 2011; Holroyd, Pakzad-Vaezi and Krigolson, 2008; Proudfit, 2015), and it is believed that the impact of positive (increase in dopamine) and negative (decrease in dopamine) RPEs on the ACC following goal-directed feedback modulates the amplitude of the reward positivity (Baker and Holroyd, 2011; Holroyd and Coles, 2002; Holroyd and Yeung, 2012). Although hotly debated, converging evidence across multiple methodologies indicate that the reward positivity reflects an RPE signal and is generated by ACC(Holroyd and Umemoto, 2016; Holroyd and Yeung, 2012).

While the reward positivity and other RPE-related EEG signals (e.g., frontal midline theta,(Cavanagh et al., 2010))have been studied for decades in highly controlled experimental task settings (e.g., participants press buttons to make choices between options that pay out probabilistic rewards), these oversimplified tasks may fail to engage real-world cognitive processes. In particular, RPE-related neural responses and behavior in these tasks has been traditionally explained with reinforcement learning models, which make quantitative predictions about observable physiological and behavioral data and, in their underlying equations, make strong claims about the unobservable algorithms used by the brain(Cavanagh et al., 2010; Doll et al., 2009; Watkins and Dayan, 1992). Yet, while the ability of these models to predict behavior in simplified lab-based experiments is firmly established, the claim that they can explain behavior in complex real-world tasks by reproducing the cognitive computations performed in the brain has proven difficult to test. Moreover, because the experiments are so simplified, it is not clear if they engage the same cognitive processes that are used in the real world.

Here, we seek to move beyond simplistic lab-based experiments by measuring RPE-related neural activity and behavior in humans while they freely perform a task in a realistic, everyday environment. To do so, we leveraged technological advances in mobile-EEG and head mounted AR glasses to investigate RPE-related processes in humans freely navigating a room to find rewards. AR is an interactive experience where virtual objects are overlayed on the real work by computer-generated perceptual information across multiple sensory modalities (e.g. visual, auditory) using a special kind of optic glasses (Fig. 1B: Hololens 2, Microsoft).

**Figure 1.**
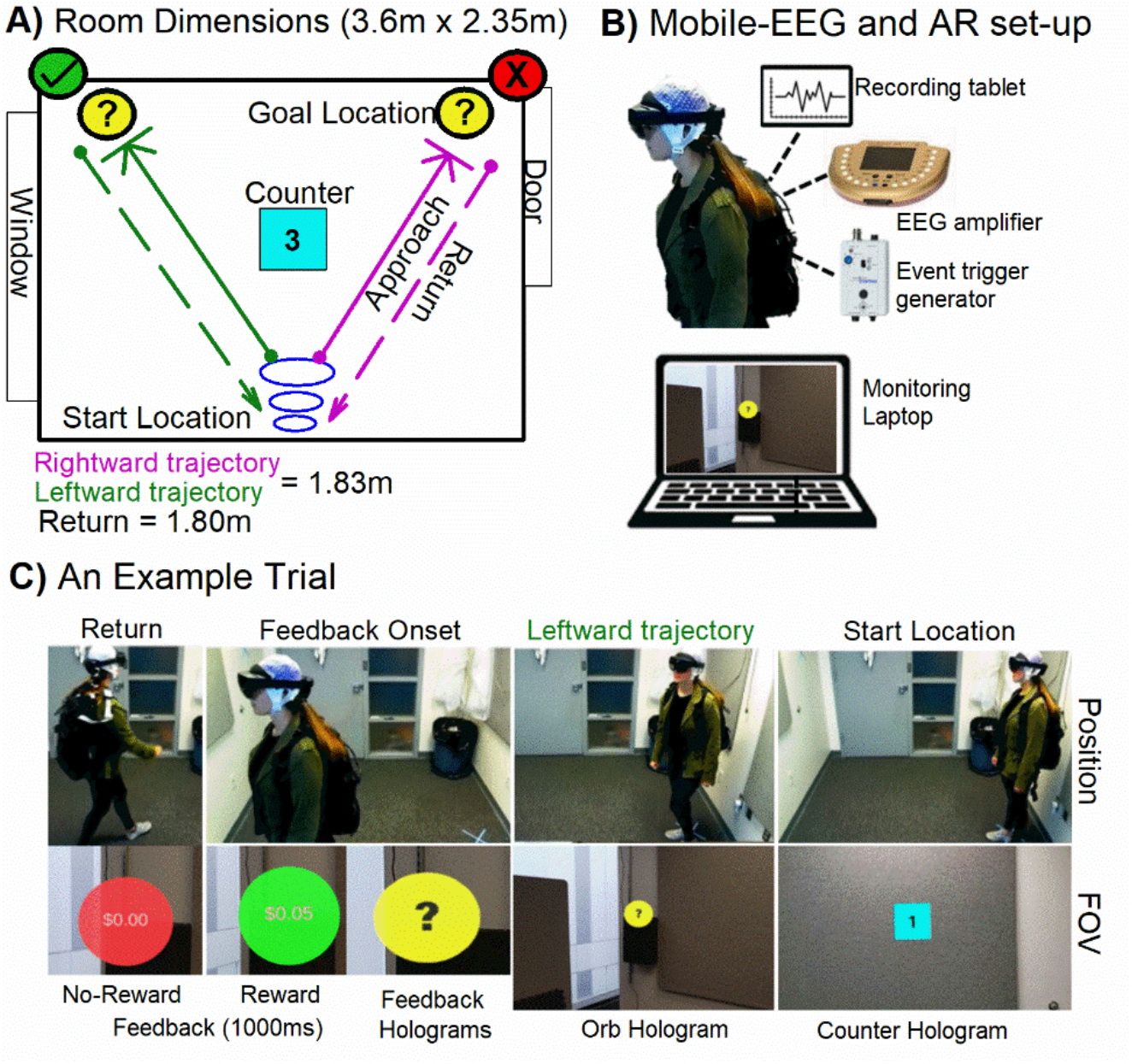
Experimental Set-up. Mobile-EEG and augmented reality (AR) operant chamber paradigm. **(A)** Dimensions of the physical room and placement of the task holograms. Purple and green lines denote rightward and leftward trajectories, respectively. **(B)** AR hardware and EEG set up, which include a Hololens2 (Microsoft), V-amp EEG system with 16-channel BrainVision actiCAP electrodes and StimTrak system to record event triggers (audio) (Brain Products GmbH, Munich, Germany), a tablet to record EEG data and a standard laptop to monitor subjects’ field-of-view (FOV) and to provide instructions. **(C)** An example of a rightward trajectory in the AR task (see SOM for a video depicts trial-to-trial sequence of events).

AR is seamlessly interwoven with the physical world such that it is perceived as an immersive aspect of the real environment. In this way, AR alters one’s ongoing perception of a real-world environment, and can therefore be an ideal solution for providing experimental control of stimulus in any real-world setting. Because the reward positivity has not yet been investigated in a real-world setting, we examined whether holographic rewards cues interwoven within the physical world could elicit this ERP component. In sum, we propose that the melding of mobile-EEG and AR provides a unique opportunity to study the ecological validity of RPE-related neural signals evoked during goal-directed behavior in humans as it holds promise for integrating experimental, computational, and theoretical analyses of laboratory behavior in a real-world setting (Krugliak and Clarke, 2021; Lange and Osinsky, 2020).

## Materials and Methods

### Participants

Twenty-five adults (24 right-handed, 6 male and 19 female, aged 18–43 years old [M = 23, SE = 6]) were recruited from the Newark community and Rutgers University Department of Psychology participant pool using the SONA system. This study was approved by the Institutional Review Board of Rutgers University and all experiments were performed in accordance with relevant guidelines and regulations. The study adhered to the principles expressed in the 1964 Declaration of Helsinki. Informed consent was obtained from all participants. Each participant received either course credit or $25 an hour plus $5 task bonus for their participation. Before the experiment, participants were screened for neurological symptoms and histories of neurological injuries (e.g., head trauma), and then asked to fill out the Edinburgh Handedness Inventory (Oldfield, 1971). After the experiment, participants filled out the Everyday Spatial Questionnaire. All participants provided written consent before the experiment.

### AR Operant Chamber

Since the operant chamber task has been the gold standard laboratory task for studying reinforcement learning and goal-directed behavior in freely moving animals (Schifani et al., 2017); (Eagle and Robbins, 2003), we utilized mobile-EEG and AR technology to create a full-scale human version of the operant chamber (Fig. 1A). The AR operant chamber was enclosed inside the lab’s physical space of a 2.13m by 2.13m room and was constructed using commercially available computer software (Unity version 2019.2, https://unity.com). The AR environment was provided through *Microsoft* Hololens2 (HL2) head-mounted holographic system, which tracked participants’ head positions and eye gaze-fixation during task performance. Continuous EEG was recorded from 16 actiCAP slim electrodes using a mobile V-Amp amplifier system (Brain Products, Munich, Germany).

Prior to the experiment, participants were trained to use the HL2 system, and performed the eye calibration set-up to ensure accurate eye tracking. Following eye calibration, participants were instructed to set-up the experiment, while the experimenter tracked their point-of-view (POV) remotely on a tablet. The set-up consisted of placing three holograms within the room, the start portal (ascending blue rings) and two yellow floating orbs marking left and right goal-locations (Fig. 1A). Participants placed the start portal over a floor sticker, and the two yellow orbs against a left and right goal-box on the back wall. The yellow orbs automatically adjusted to the participants’ eye level. Once the holograms were placed in the correct locations, participants were instructed to stand in the start portal and to press a virtual button to begin the experiment. Once standing in the start location, a holographic countdown timer would appear for 3 seconds, and participants were instructed to make their choice at the end of the countdown. Depending on their choice, participants could move towards the left or right goal-location (yellow floating orb), and once standing in front of the orb and looking at it as detected using eye tracking, the floating yellow orb turned either green to signify a reward (5 cents) or red signifying no-reward (0 cents) (Fig. 1C). Following the feedback (duration 1000 msec), participants then walked back to the start location to begin the next trial. The task consisted of four blocks (50 trials per block), separated by self-timed rest breaks that presented their cumulative earnings. Unknown to them, on each trial the type of feedback was selected at random (50% probability for each feedback type), a necessity to record the reward positivity using a difference wave approach (Cockburn and Holroyd, 2018). At the end of the experiment, participants were informed about the probabilities and were given a $5 performance bonus.

### HL2 data acquisition

The software running on HL2 recorded and exported experimental and behavioral data (e.g., timing, position, distance, eye-gaze, feedback, tone onset/offset and countdown; 1=event, 0=nonevent) for each participant (Fig. 1A). The sampling rate of the HL2 was 60hz and recorded via Bluetooth to a comma-separated values file on a remote computer in the adjacent room. Time-locked EEG markers were sent to the EEG system (Fig. 1B) by converting an 11 msec event-related audio signal or sine wave (e.g., countdown onset and offset, feedback onset and offset) to a TTL pulse using the BrainVision (BV) StimTrak system (Gain factor = 1; Trigger Level= 3V) (Fig. 1B). To note, there was an unexpected delay (< 200 ms) between visual and auditory onset which could not be corrected in the HL2 programming platform, which resulted in a delay of the event triggers recorded in the EEG system. To correct for the output-input delay, the HL2 event markers and EEG triggers were synchronized using custom written MATLAB scripts that added 200ms to each trial. HL2 activity was monitored and controlled using both the web browser access (*Microsoft Device Portal*) and the HoloLens Application (*Microsoft Corporation*, version 1.1.70) on an external Microsoft tablet and laptop.

### Electrophysiological Data Recording

The electroencephalogram (EEG) was collected using a 16-channel actiCAP snap system (Brain Products GmbH, Munich, Germany) with 12 scalp electrode sites (C3, C4, Cz, F3, F4, FC1, FC2, FCz, P3, P4, P7, and P8) and four external electrodes. The EEG signals were referenced online to channel Pz with a ground at AFz, amplified using a portable V-Amplifier, and recorded using Brain Vision Recorder software (Brain Products GmbH, Munich, Germany). The sampling rate was set to 1000 Hz. The electroocculogram (EOG) was recorded for the purpose of eye-artifact correction. Horizontal EOG was recorded from the external canthi of both eyes, and vertical EOG was recorded from the suborbital and infraorbital regions of the right eye. To note, by convention mastoid sites (M1 and M2) were collected to re-reference offline. However, these electrodes were removed from the dataset due to excessive noise and were not used in the analysis (Lin et al., 2022).

### Electrophysiological Data Analysis

EEG data were analyzed offline using BrainVision Analyzer 2 (Brain Products GmbH, Munich, Germany). The EEG signals were filtered using a fourth-order digital Butterworth filter with a bandpass of 1-20 hz. Eye-artifacts were corrected using independent component analysis method (ICA) and inverse component analysis which splits the signals up into independent eye components, removes the component from the EEG, and projects it back into the time-domain (Jung et al., 2000). The EEG data were then segmented into 1000 msec epochs spanning from -200 to 800 msec from feedback onset. The segmented data was then baseline-corrected using a mean voltage range from -200ms to 0ms. The data was then re-referenced using an average reference created from the following channels: C3, C4, Cz, F3, F4, FC1, FC2, FCz, P3, P4, P7, and P8. Segments containing muscular and other artifacts were removed using the following criteria: (1) a maximal voltage step of 50μV/msec, (2) a maximal difference of values in intervals of 250μV, and (3) lowest allowed activity values in intervals of 0.5μV. Following artifact rejection, channels containing artifacts that exceed 5% of the data were identified and interpolated using Hjorth-nearest neighbor algorithm (Hjorth, 1975). Prior to averaging, we corrected the latency jitter in the ERP across trials by applying the Adaptive Woody Filter method (AWF) using a 100-300ms time-window with 50ms step interval at channel FCz (Li et al., 2009; Lin and Baker, 2022; Woody, 2006).

ERPs were created for each participant and electrode by averaging the single-trial EEG data according to feedback type (reward and no-reward feedback). The reward positivity was then evaluated as a difference wave by subtracting reward from no-reward ERPs. The size of the reward positivity was then determined by identifying the peak amplitude of the difference between the reward and no-reward ERPs within a 100-to 400-msec window after feedback onset. The difference wave method was recommended in a meta-analysis and isolates the reward positivity from other ERP components (Sambrook and Goslin, 2015). Local maxima peak detection was used on the difference wave to extract peak amplitude of the reward positivity at each channel within the 100 to 400ms time-window (Baker et al., 2020; Baker et al., 2016; Baker, Wood and Holroyd, 2016; Biernacki, Lin and Baker, 2020). To confirm topographical distribution of the reward positivity, a paired sample t-test was performed between the reward positivity maximal (channel FCz) compared to all other channels. Peak amplitudes, at each channel, were also tested against zero using one-sample t-test with a significance level of a = 0.01 to confirm the presence of the reward positivity.

### Behavioral Analysis

Operant chamber performance measures included 1) reaction time (RT) reported in seconds and measured from count-down offset to feedback onset (start RT), and from feedback location to the start location (return RT); 2) post-feedback RT measured from feedback onset back to start location; and 3) win-stay and lose-shift behavior; defined by choosing the same location (right or left) after a reward feedback and selecting the alternative location after no reward feedback, respectively. We excluded trials with RTs slower than 5% of the higher boundary. Post-feedback performance was analyzed using general linear models (GLM) with feedback (win/loss), behavior (stay/shift) and task-half (Half 1: Block 1-2, 1-99 trials; Half 2: Block 3-4, 100-200 trials) as within-subject factors, followed by post-hoc tests with a significance level of α = 0.05. Out of 25 participants, data from 4 participants were excluded from final analyses due to: multiple bad channels (n = 1), protocol error (n = 1), marker file mismatch (n = 1), and EEG recorder technical failure (n = 1).

## Results

### Task Performance

Overall, participants completed 195 trials (SD = 14, range = 150 – 200), and took about 3.38 seconds (±0.93s) to reach the feedback location (∼2.49m). A two-way repeated measures ANOVA on start-RT with Half (Half-1, Half-2) and Direction (Left vs. Right) as factors revealed a main effect of Half, F(1, 21) = 6.35, p <0.05, η^2^ = 0.23. Post-hoc tests indicated a faster RT for the second half (M=2.39s ± 0.10s) compared to the first half (M = 3.40s ± 0.11s), t (21) = 8.87, p<0.001, Cohen’s D= 1.86, (Fig. 2A). Further, there was a main effect of direction, F (1, 21) = 4.33, p < 0.05, η^2^ = 0.17, showing that participant approached the right target (M=3.29s ± 0.10s) slightly faster than the left target (M=3.43s ± 0.10s), t (21) = -1.95, p=0.06, Cohen’s D = 0.42. Regarding choice behavior, no main effects nor interactions were found.

**Figure 2.**
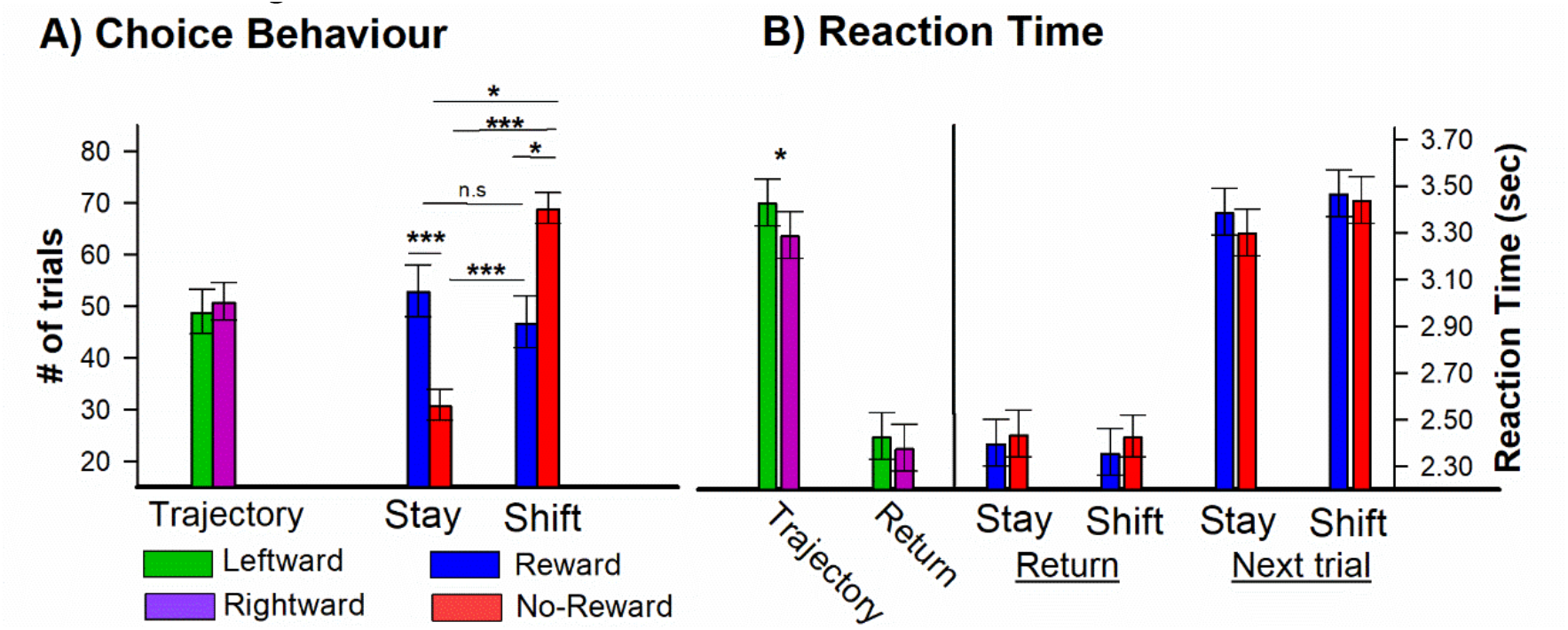
Results of the AR Operant Chamber performance analysis. Behavioral analysis for choice (A) and reaction time (B). Green and purple bars denote leftward and rightward trajectories, and Blue (reward) and Red (no-reward) bars denote post-feedback return behavior (feedback location to start-location) and next trial behavior (start location to feedback). To note, although not shown here, reaction time was slower during the return to start-location following no-reward cues (M=2.43s ± 0.10s) compared to reward cues (M= 2.38s ± 0.11s), p<0.05. Significant effects are shown as follows: *p < .05, **p < .01, ***p < .001, (two-tailed). Error bars denote standard error.

Next, a repeated measures ANOVA on post-feedback choice with feedback (win vs. loss), behavior (stay vs. shift) and task-half (Half 1 vs. Half 2) as within-subject factors revealed a main effect of behavior, [F (1, 21) = 4.31, p < .05, η^2^ = .17. Post hoc analysis indicated that participants shifted (58% ± 3.8) more often than stayed (42% ± 3.8), t (21) = 2.08, p<0.05, Cohen’s D = 0.81. This analysis also revealed an interaction between feedback and behavior, F (1, 21) = 15.38, p < .001, η^2^ = .42. Post-hoc test indicated that participants shifted their responses more often following negative feedback (M=68.3% ± 3.5) compared to positive feedback (M= 47.1%, ± 5.7), t (21) = 3.67, p<0.01, Cohen’s D =0.95 (FIG. 2B). By contrast, participants stayed with their response more often following positive feedback (M=52.8%, ± 5.7) compared to negative feedback (M= 31.7%, ± 3.5), t (21) = 3.67, p<0.01, Cohen’s D=0.95 (FIG 2B). To note, no differences were observed between win-stay and win-shift performance (p >.05), but a difference was observed between lose-shift and lose-stay (p<.001). Even though feedback was randomized, these results indicated that feedback influenced the participants’ subsequent behavior in this task. In regards to post-feedback RT, there was a main effect of Half, F(1, 21) = 4.23, p < 0.05, η^2^ = 0.18. indicating a faster RT for the second half (M=2.39s ± 0.11s) compared to the first half (M = 3.45s ± 0.10s). Finally, a repeated measures ANOVA on post-feedback RT during the return with feedback (win vs. loss) and behavior (stay vs. shift) as within-subject factors revealed a significant main effect of feedback, F (1, 21) = 4.02, p < 0.05, η^2^ = .16, indicating that RT was slower following no-reward cues (M=2.43s ± 0.10s) compared to reward cues (M= 2.38s ± 0.11s), t (21) = -2.01, p<0.05, Cohen’s D = 0.42 (Fig. 2C). No other main effects nor interactions were observed.

### Reward Positivity

Because the reward positivity to holographic feedback stimuli has not yet been investigated, we first examined whether holographic rewards encountered during free navigation elicit this ERP component. Fig. 3 presents stimulus-locked grand averages for both feedback conditions at channel FCz. Consistent with previous research, the reward positivity elicited by monetary rewards was clearly evident in the difference wave (M= -2.60μV, ± 1.73μV) peaking 223 ms after feedback onset (Fig. 3A, solid lines), and was significantly different from zero, t (20) = -6.88, p < 0.01, Cohen’s D= -1.5). Further, as shown in Fig. 3A, the frontal–central distribution is consistent with the identification of this ERP component as the reward positivity (Miltner, Braun and Coles, 1997), indicating that holographic-related feedback can elicit the reward positivity in freely moving participants. These results confirm that the AR operant chamber paradigm elicited the reward positivity component. To note, this paradigm also elicited other common feedback-related ERP components, particularly the N100, P100, N170, and P200 (see Fig. 3B), indicating that the AR task is capable of eliciting a broad spectrum of cognitive processes, including those related to perceptual processing, cognitive control, and contextual updating (Luck, 2014).

**Figure 3.**
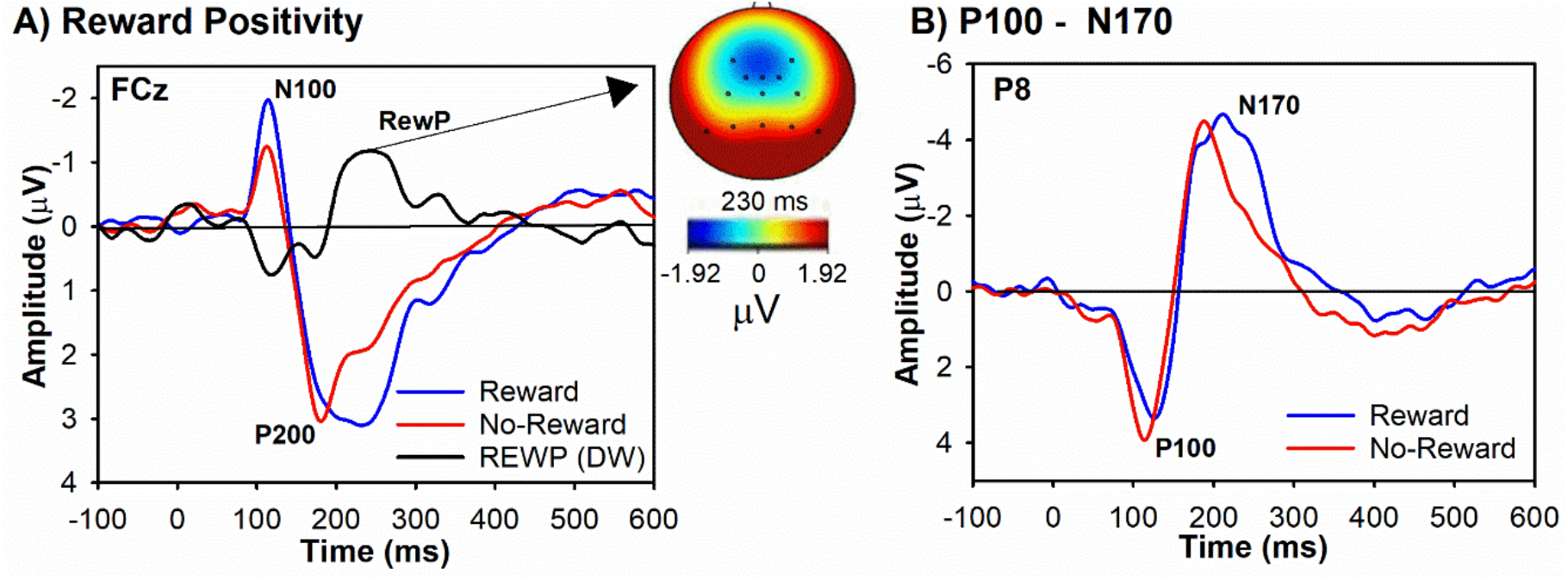
Reward Positivity results. A) ERPs elicited by reward feedback (blue), no-reward feedback (red) and difference wave (black-reward positivity) averaged across all blocks. Topoplots denote the amplitude of the reward positivity at 230 msec. B) For the purpose of comparison, we plotted the ERPs over posterior channel P8 to highlight that the feedback stimulus presented in the AR paradigm is capable of eliciting ERPs commonly associated with perceptual processing of the stimulus (N100 and N170). Data are associated with channel FCz (left panel) and P8 (right panel) and negative is plotted up by convention.

## DISCUSSION

While the electrophysiological response to rewards has been well documented during laboratory-based tasks, little is known about the ecological validity of these responses in ‘real-world’ settings. To examine this issue, we designed a novel AR operant chamber task to test whether holographic reward cues presented in the real-world can elicit the reward positivity, an ERP component associated with ACC sensitivity to RPE signals. To the best of our knowledge, we are the first to show that the reward positivity can be accurately recorded during real-world behavior, and that participants performed the task in accordance with reinforcement learning theory. Foremost, the reward positivity peaked at around 235ms post-feedback with a frontocentral negative topographic distribution, replicating previous reward positivity studies using tasks displayed on desktop computers (Baker and Holroyd, 2009, 2011; Baker et al., 2016; Proudfit, 2015; Sambrook and Goslin, 2015). An influential theory of ACC function proposes that the ACC utilizes dopaminergic RPEs signals to learn the value of rewards for the purpose of selecting and motivating the most appropriate action plan directed toward goals(Holroyd and Coles, 2002; Holroyd and Umemoto, 2016; Holroyd and Yeung, 2012). Accordingly, it is believed that the impact of positive RPEs on the ACC following goal-directed feedback modulates the amplitude of the reward positivity(Baker and Holroyd, 2011; Holroyd, Pakzad-Vaezi and Krigolson, 2008; Holroyd and Umemoto, 2016). Converging evidence across multiple methodologies indicate that the reward positivity reflects an RPE signal and is generated by ACC (Holroyd and Umemoto, 2016; Holroyd and Yeung, 2012; Sambrook and Goslin, 2015). In particular, genetic, pharmacological, and neuropsychological evidence implicates dopamine in reward positivity production (Baker et al., 2016); and source localization studies, simultaneous recording of electroencephalogram/functional magnetic resonance imaging (EEG/fMRI) data, and intracranial recording studies in rodents (Warren et al., 2015), nonhuman primates (Emeric et al., 2008), and humans (Ramakrishnan, Hayden and Platt, 2019) indicate that the reward positivity is generated in ACC. Taken together, these results indicated that the AR-version of an operant chamber task is capable of eliciting RPE-related ACC activity in a real-world setting, as revealed by the observation of the reward positivity.

At a behavioral level, participants exhibited a lose-switch strategy and walked slower from the goal location to the start location following no-reward feedback, evidence that the AR task can drive adaptive learning. More specifically, participants shifted more often following negative feedback compared to positive feedback and repeated their response more often following positive feedback compared to negative feedback. These results indicate that feedback did in fact influence behavior and appears consistent with Thorndike’s Law of Effect: if an action is followed by a reward or punishment then that action will be more or less likely, respectively, to reoccur (Catania, 1999). Further, given that RPE signals are used for the purpose of action selection and reinforcement learning, these results could reflect the degree in which negative and positive RPEs modify behavior. For example, we observed a trend towards post-error slowing while subjects walked from the goal-location to the start location. Post-error slowing represents the amount that responses slowed on a trial following an erroneous behavioral response (or negative feedback) compared to a trial following a correct response (or positive feedback) (Schroder et al., 2020); (Heydari and Holroyd, 2016). Varying accounts suggest that post-error slowing is reflective of the degree RPE signal are utilized for future behavioral adaption (immediate reaction time slowing following errors), fitting with the proposed function of the prefrontal cortex and ACC in cognitive control and reinforcement learning (Dutilh et al., 2012; Yeung, Botvinick and Cohen, 2004). Although a trend, it is worth noting that this is the first-time post-error slowing has been observed beyond button press tasks.

These results add a compelling dimension to our understanding of how rewards shape decision-making and learning. While we found that participants maintained their chosen behavior following reward feedback compared to no-reward feedback, it is noteworthy that they exhibited an equal propensity to modify their response after a reward, demonstrating no difference between win-stay and win-shift performance. This observation deviates from heuristic learning models typically employed in various fields, including psychology, game theory, statistics, economics, and machine learning, where the win-stay strategy often dominates. However, this is the second time we replicated this result, previously reported similar findings in a study where participants navigated an immersive virtual reality T-maze task to locate rewards (Lin et al., 2022). Conversely, in traditional experiments where subjects are merely pressing buttons to make decisions on a computer screen, a higher proportion of win-stay responses compared to win-shift responses is observed (Baker et al., 2020; Baker, Wood and Holroyd, 2016). One possible explanation for this discrepancy might lie in the differential cognitive demands between active navigation and simple button-press tasks (Coddington and Dudman, 2019). During active navigation, participants may be more inclined to explore various strategies (i.e., hypothesis testing through frequent win-shift behavior) due to the heightened cognitive and physical effort required to navigate their bodies towards a goal. In contrast, button-press tasks, which demand minimal physical or cognitive effort, might promote more conservative, win-stay behavior. While these results require further empirical testing, they do present compelling evidence that participants making decisions between options presented in a real-world setting may computationally differ from participants pressing buttons in simplistic lab-based experiments. Thus, the ability of current reinforcement learning models to predict behavior in simplistic lab-based experiments may be insufficient for explaining behavior and cognition in the complexity of real-world tasks and are ripe for future investigations.

Taken together, the observation of the reward positivity and adaptive behavior in this real-world task strengthens the ecological validity (EV) of goal-directed behavior. EV refers to three dimensions of experimentation (research setting, stimuli, and response) that should mimic the natural world as closely as possible (Brunswik, 1943; Lewin, 1943; Schmuckler, 2001). Research setting EV concerns the environment in which the research takes place; stimuli EV addresses the issue of representativeness and naturalness of objects presented in an experiment; and response EV involves the nature of the task and behavior required from the participant (Brunswik, 1943; Schmuckler, 2001). The current study addresses the three EV dimensions of goal-directed behavior by: 1) utilizing mobile-EEG and AR methods to create a real-world operant chamber (*research setting EV*), 2) demonstrating the ability to record ACC-related electrophysiological responses (e.g., the reward positivity) to holographic reward cues in the real-word (*stimuli EV*), and 3) revealing adaptive responding based on positive and negative feedback (*response EV*). Regarding research setting EV, AR provides an opportunity to interweave and control experimental manipulations within the participants physical world, thereby altering their ongoing perception of real-world events. While previous studies have used virtual reality environments to balance naturalistic observations and control (Campbell et al., 2009; Parsons, 2015), inherent limitations emerge – motion sickness, limited the range of navigation, computer-generated environments do not truly reflect “our world”, the inability to see their own bodies creates a sense of disembodiment, and extensive training (Garrett et al., 2018). AR methods overcome these limitations, thereby providing an exciting opportunity to conduct future experiments beyond the laboratory and in more natural settings. In relation to stimuli EV, we found that reward-related holograms can elicit the reward positivity. As argued elsewhere, replication of neural signatures found in desktop versions of a similar real-world task demonstrate the feasibility of such approaches while simultaneously accounting for EV of the task in general (Krugliak and Clarke, 2021). In one notable instance, Lange & Osinsky (2020) were able to replicate an increased frontal-midline theta responses to negative action outcomes during a real-world toy shooting task but failed to elicit feedback-related ERPs, likely due to the sensitivity of ERPs to stimulus onset (i.e., latency jitter) (Lange and Osinsky, 2020). Finally, regarding response EV, the AR task drove adaptive learning following negative feedback but failed to elicit a dominant win-stay strategy. Collectively, these findings underscore the promise of integrating mobile-EEG and AR to enhance the EV of electrophysiological studies of goal-directed behavior. This methodological innovation broadens the horizons for conducting human cognitive neuroscience research, moving beyond laboratory confines and into more ecologically valid settings.

## CONCLUSION

In sum, the present study provides evidence that combining mobile-EEG with AR technology is a feasible solution to enhance the ecological validity of human electrophysiological studies of reinforcement learning and goal-directed behavior and holds promise for integrating experimental, computational, and theoretical analyses of goal-directed behavior in animals within the field of human EEG research. Future clinical applications for this paradigm could also uncover open questions of cue reactivity in drug addiction and other mental health disorders.

## Limitations

Although this research presents the first data using mobile-EEG and AR to examine electrophysiological and behavioral response properties during active goal-directed navigation, future research may address some of the study’s limitations. First, while the ERP results were consistent with previous work, the amplitude of the reward positivity appeared smaller compared to the reward positivity recorded from stationary subjects in a highly controlled environments (Baker and Holroyd, 2009; Baker, Wood and Holroyd, 2016). While undoubtfully active walking decreased the signal-to-noise ratio, and thus reduced the amplitude of the reward positivity and other feedback-related ERPs (e.g., P200), it is worth noting that the amplitude of other ERPs associated with sensation and perception (P100 and N170) were consistent with previous work (Baker and Holroyd, 2013). Future studies should attempt to dissociate whether the small amplitudes observed here was a result of methodological confounds (i.e., device-related trial-to-trial latency jitter may distort ERP amplitudes) or a natural phenomenon observed during complex real-world tasks (i.e., cognitive effort/energy may be allocated or distributed across multiple systems). Another limitation inherent to testing EV is that the more the environment is naturalistic, the less it lends itself to experimental control. Any natural phenomena existing during our day-to-day activity (e.g., weather, bystander interference, overstimulation) could affect the cognitive computations performed during a natural dynamic experiment. Since the present study was still conducted within the laboratory, the current AR task may not fully represent a true real-world setting. Hence, future experiments could test this paradigm in a more dynamic world full of distractions and noise. Finally, we did not test the EV of reinforcement learning models in this proof of concept study. Future studies should test whether conventional computational models of cognition can make quantitative predictions about observable behavior in both simplistic lab-based experiments and in complex real-world tasks (Robles et al., 2021).

## Acknowledgments

We are grateful to the research assistants of the Laboratory for Cognitive Neuroimaging and Stimulation for help with data collection.

## Funding

This research was supported by Rutgers Research Council Grant and departmental research start-up funds from Rutgers University (to TEB). JS was supported by the National Institutes of Health NIGMS 5T32GM140951.

### Author contributions

Conceptualization: TEB, OL

Methodology: TEB, OL, ML, JS

Investigation: JS, TEB, ML

Visualization: TEB

Supervision: TEB

Writing—original draft: TEB, JS

Writing—review & editing: OL, ML, TEB

## Data and materials availability

The data can be provided by JS pending scientific review and a completed material transfer agreement. Requests for the data should be submitted to.

## SUPPLEMENTARY

### ERP components

A consideration of AR replication is in interpreting additional ERP components in the post-feedback waveform, particularly the P2, N2, and P3. The P2 was measured using a local peak detection method by identifying the maximum positive value of the ERP within a window extending from 100ms to 250ms following the presentation of the feedback, the time of which was taken as the peak latency of the P2. This algorithm was applied to the Reward and No-reward ERPs associated with electrode site FCz. The N2 was also quantified at FCz, and its amplitude was measured for each participant and feedback condition (reward, no reward) using a local peak detection algorithm, in which the most negative value within a time window starting from 200ms to 400ms after feedback presentation was taken as the peak. Finally, the P3 amplitude was measured by identifying the maximum positive-going value of the Reward and No-reward ERPs recorded at electrode site Pz, within a window from 300 to 600ms post-feedback.

Statistical and visual inspection revealed significant peak amplitude differences between the reward (M=0.38μV, ± 0.50μV) and no reward (M=-0.72μV, ± 0.41μV) N2 components at FCz (Fig. 2A), (t (20) = 2.79, p<0.01; *Cohen’s D*= 1.81) (Fig. 3). These results strengthen the hypothesis that there’s a dampening of the N2 by the Reward Positivity for rewarding conditions (Baker and Holroyd, 2011; Holroyd, Pakzad-Vaezi and Krigolson, 2008; Holroyd and Umemoto, 2016). No significant difference between reward and no reward were detected for the P2 and P3 (P>.05). The results suggest the Reward Positivity affects the N2 component but not the P2 and P3 in the reward condition.

